# HIV-1 Vpr counteracts TASOR restriction to promote infection prior to integration

**DOI:** 10.64898/2026.03.06.710055

**Authors:** Joseph M. Gibbons, Kelly M. Marno, Corinna Pade, Wing-Yiu Jason Lee, Áine McKnight

## Abstract

HIV-1 is challenged by many intrinsic antiviral factors including APOBEC3G, Tetherin/BST-2 and SERINC5 which are overcome by viral accessory proteins Vif, Vpu and Nef respectively. HIV-1 Vpr has multiple proposed cellular targets including REAF, previously shown by us to restrict early after virus entry, but its function is incompletely understood. TASOR is a component of HuSH, an epigenetic silencing complex which inhibits expression of HIV-1 and is overcome by HIV-2/SIV Vpx. We report that TASOR, in addition to silencing integrated provirus, inhibits viral replication prior to integration during reverse transcription. TASOR impairs viral replication in primary macrophages and restriction of a primary isolate is also shown. The restriction to replication is overcome by HIV-1 Vpr which induces TASOR degradation concomitant with reverse transcription. Using Vpr mutants Q65R and F34I we show that interaction with the CRL4-DCAF1 E3 ligase and nuclear localisation are required for Vpr to mitigate TASOR. The pre and post integration mechanism of Vpr mediated TASOR mitigation is genetically segregated using cell cycle Vpr mutant R80A. TASOR depletion induces cells to accumulate in G2/M and arrested cells are more susceptible to HIV-1. Understanding the mechanistic details of pathogen-host interactions is critical for the design of effective anti-HIV/AIDS therapeutics.

## Introduction

Upon infection of human cells, viruses navigate a network of constitutively expressed antiviral factors that challenge every stage of viral replication. These molecules both directly inhibit viral replication and mediate downstream innate and adaptive immune responses ^1^. In response, viruses have evolved to subvert the activities of these molecules. Viral accessory proteins that were once thought to be dispensable are now understood to be critical for replication of the virus *in vivo* and one of their roles is to specifically counteract restriction factors in target cells ^2,3^. Human transgene activation suppressor (TASOR/C3ORF63/FAM208A) and RNA-associated early-stage antiviral factor (REAF/RPRD2) are two such antiviral genes that were identified in a whole genome siRNA screen for restriction factors to human immunodeficiency virus 1 (HIV-1) ^4^. Previously we showed that REAF restricts the efficiency of reverse transcription of the viral genome and that upon entry into the cell, HIV-1 accessory protein Vpr antagonises REAF and promotes its degradation via the cellular proteasome ^5,6^. In a non-lethal forward genetic screen in near-haploid KBM7 cells, TASOR was shown to be involved in epigenetic repression of transcription by directing assembly of the human silencing hub (HuSH), consisting also of M-phase phosphoprotein 8 (MPP8) and Periphilin (PPHLN1)^7^. HuSH is an RNA-mediated transcriptional silencing system that associates with RNA processing components, protects the genome from foreign genetic material and also plays a role in immune signalling ^8,9^. The HuSH subunits TASOR, MPP8 and PPHLN1 act together to mediate transcriptional silencing at genomic loci rich in H3K9me3 including newly integrated retroviruses by both reading and writing of H3K9 methylation via recruitment of the H3K9 methyltransferase SET domain bifurcated 1 (SETDB1) and the ATPase MORC2 ^7,8,10–12^. HuSH is also recruited by NP220/ZNF638 to silence unintegrated murine leukaemia virus (MLV) DNA but TASOR depletion does not give rise to elevated expression from unintegrated HIV-1 DNA ^13^. A recent analysis of TASOR interactions showed that it also interacts with a CNOT1 RNA degradation pathway as well as RNA Pol II ^14^.

Human Immunodeficiency Virus 2 (HIV-2) and Simian Immunodeficiency Virus (SIV) Vpx and Vpr from specific SIV lineages can induce both the down-modulation of TASOR and reactivation of a latent transcriptionally silent HIV-1 LTR reporter genome ^15,16^. Like many viral accessory genes, they are involved in the counter restriction of multiple restriction factors ^17^. In addition to mediating TASOR inhibition, Vpx also counteracts restrictions involving SAMHD1 and a SAMHD1-independent block to reverse transcription ^18–20^. Vpx/Vpr antagonism of HuSH and SAMHD1 demonstrates its molecular plasticity and it is also host species-specific. For example, TASOR is downmodulated in AGM VERO cells but not in owl monkey OMK cells by Vpx from HIV-2/SIVsmm ^21^. HIV-1 Vpr and HIV-2/SIV Vpx are paralogs with sequence homology and are unique amongst the accessory proteins in that they are actively incorporated into virus particles via the p6 domain of Gag ^22–25^. This suggests an important activity early in infection prior to integration of the viral genome when *de novo* Vpr can be produced. HIV-1 Vpr has multiple pleiotropic effects in the infected cell and contributes to uncoating, reverse transcription, nuclear import and transcription events amongst others ^26–29^. It is particularly important for replication of the virus in macrophages and alters both the transcriptional and proteomic landscape of the infected cell post infection ^30–36^.

Vpr targets multiple specific cellular proteins, including REAF ^6^, the structure-selective endonuclease MUS81-EME1^37^, helicase-like transcription factor (HLTF) ^38^ and Uracil-DNA glycosylase-2 (UNG2)^39^ for degradation via the proteasome using CRL4 E3 ubiquitin ligase complex and its cellular partner DCAF1. Lysosomal-associated transmembrane protein 5 (LAPTM5), which decreases progeny virion infectivity by redirecting viral Env for lysosomal degradation, is also down-modulated by HIV-1 Vpr^40^. While enhanced replication kinetics of HIV-1 are seen in the absence of TASOR, some groups show minimal effects of HIV-1 Vpr on TASOR protein levels and the precise relationship between TASOR, HIV-1 replication and Vpr remains elusive^4,9,13,15^. Others show that HIV-1 Vpr prevents the phosphorylation of TASOR (T819) early in mitosis, a process distinct from HIV-1 reactivation upon TASOR knock down ^41^. Laguette *et al.* describe the interaction between Vpr and SLX4 which leads to the recruitment and degradation of MUS81 and premature activation of the SLX4 complex resulting in cell cycle arrest in G2/M ^37^. They also found that TASOR (there called C3ORF63), along with MUS81 and UNG2, co-purified with Vpr from THP-1 cells ^37^. While it is clear that HIV-1 Vpr is responsible for cell cycle arrest at G2/M upon infection, the precise mechanism and advantage to the virus are not clear ^42–44^.

Here we report an analysis of the relationship between TASOR and HIV-1 Vpr in susceptible cells where Vpr enhances replication efficiency. We show that TASOR activity restricts a primary isolate of HIV-1 and impairs reverse transcription of the viral genome. HIV-1 Vpr overcomes TASOR mediated restriction and can deplete TASOR in primary macrophages, making them more susceptible to infection. TASOR protein is rapidly lost early after viral infection and this is dependent on Vpr. Depletion of TASOR results in G2/M arrest and arrested cells are more susceptible to infection. Our data is a further demonstration of the multifaceted nature of HIV-1 Vpr and highlights its attractiveness for the development of novel therapeutics. It also identifies a novel restrictive activity of TASOR prior to integration of the viral genome. Understanding host-pathogen interactions is critical for targeted drug design against HIV/AIDS.

## Results

### HIV-1 Vpr overcomes TASOR mediated restriction and promotes infection in macrophages

A critical role of viral accessory proteins is to specifically target and downmodulate antiviral proteins to overcome their restrictive effects ^17^. The abilities of SIV Vpx to downmodulate and challenge TASOR activity and HIV-1 Vpr to downmodulate and challenge REAF have been previously documented ^6,7,15,16^. We show by Western blotting of whole cell lysates from Jurkat cells that SIV Vpx or HIV-1 Vpr alone are sufficient to induce to downmodulate both REAF and TASOR (Figure 1a). Delivery of HIV-1 Vpr, and to a much lesser extent SIV Vpx, also induced an accumulation of cells in the mitotic phase, as indicated by the elevated levels of phosphorylated histone H3 (Ser10/Thr11).

**Figure 1:**
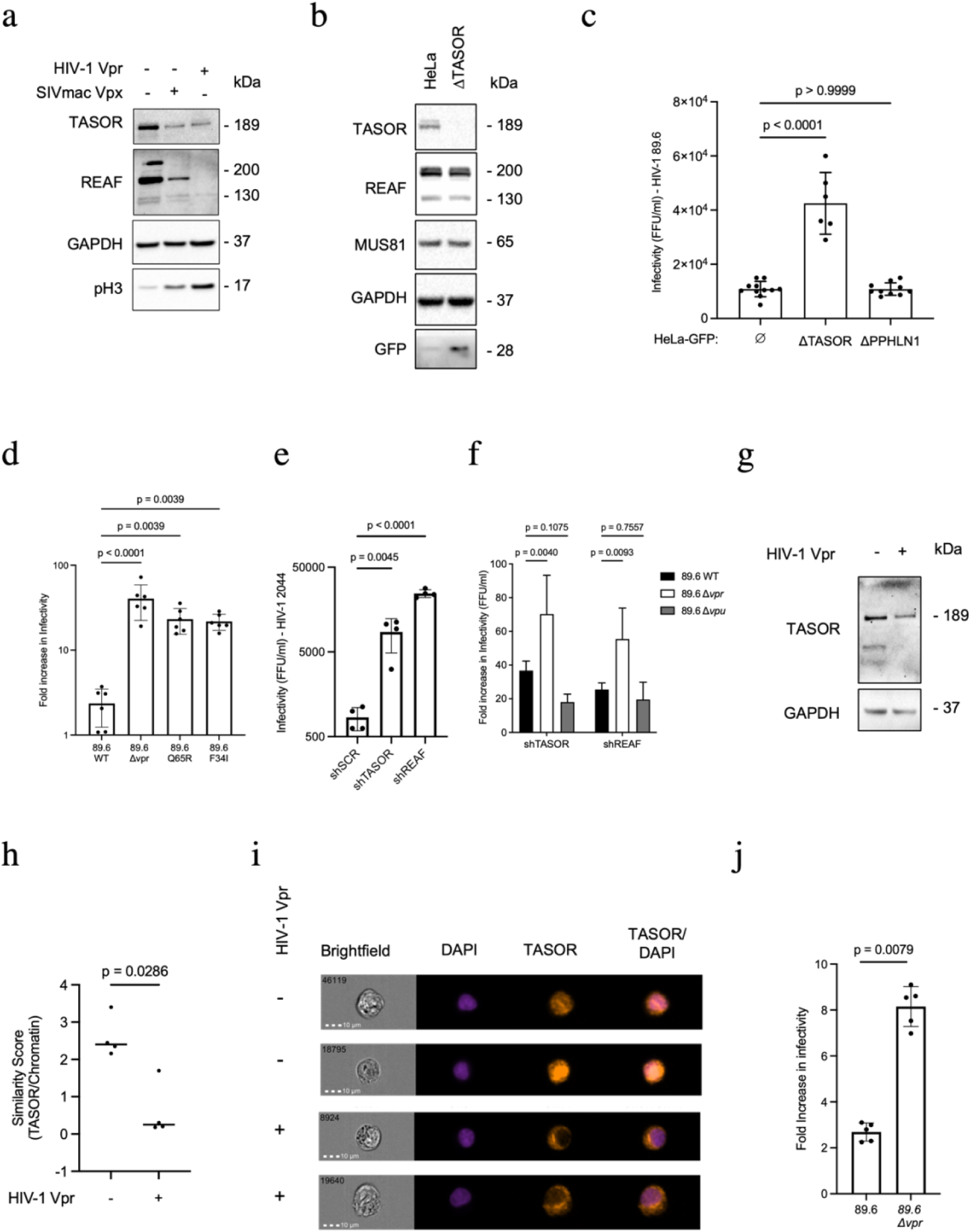
HIV-1 Vpr overcomes TASOR mediated restriction and promotes infection in MDMs. **a**, Western blotting of TASOR, REAF and phosphorylated histone H3 (pH3 Ser10/Thr11) in Jurkat cells treated for 24 hours with HIV-1 Vpr or SIVmac239 Vpx VLPs (50ng measured by p24 ELISA). **b**, Western blotting of REAF, TASOR, MUS81 and GFP in HeLa and HeLa-ΔTASOR. **c**, Infectivity (FFU/ml) of HIV-1 89.6^WT^ on HeLa-CD4 (∅), HeLa-ΔTASOR-CD4 and HeLa-ΔPPHN1-CD4. Error bars indicate standard deviations of the means derived from duplicate titrations. Ordinary one-way ANOVA with multiple comparisons and Dunnet’s test. **d**, HeLa-CD4 and HeLa-ΔTASOR-CD4 challenged with HIV-1 89.6^WT^, HIV-1 89.6^Δ*vpr*^, HIV-1 89.6^Q65R^ or HIV-1 89.6^F34I^. Fold increase in infectivity measured by ELISA of p24 protein in cell culture supernatant. Error bars represent standard deviations from the means of replicates. Ordinary one-way ANOVA with multiple comparisons and Holm-Šídák correction. **e**, Infectivity (FFU/ml) of HIV-1 2044 on HeLa-shTASOR-CD4 and HeLa-shREAF-CD4. Error bars represent standard deviations from the means of titrations performed in duplicate. Ordinary one-way ANOVA with multiple comparisons and Dunnet’s test. **f**, Fold increase in infectivity (FFU/ml) of HIV-1 89.6^WT^, HIV-1 89.6^Δ*vpr*^ and HIV-1 89.6^Δ*vpu*^ on HeLa-CD4 cells with shRNA targeting TASOR/REAF compared to control shRNA cells. Error bars represent standard deviations from the means of titrations performed in duplicate. Two-way ANOVA with multiple comparisons and Dunnet’s correction. **g**, Western blotting of TASOR in primary MDMs treated for 24 hours with VLPs containing HIV-1 Vpr or control VLPs. **h**, Imaging flow cytometry of TASOR/Chromatin Similarity Score (co-localisation) in DAPI stained MDMs treated for 24 hours with VLPs. Mann-Whitney t-test. **i**, Representative images (60X magnification) of VLP treated MDMs from g and h which were treated Vpr-containing or empty VLPs. **j**, Fold increase in infectivity (p24 production over 48 hours) for HIV-1 89.6^WT^ and HIV-1 89.6^Δ*vpr*^ infected MDMs previously treated with HIV-1 Vpr containing VLPs relative to empty control VLP treated MDMs. Error bars represent standard deviations from the means of replicates. Mann-Whitney t test.

HeLa-GFP cells contain a GFP reporter under HuSH mediated repression ^7^. CRISPR-Cas9 knock out of TASOR (HeLa-GFP-ΔTASOR) results in de-repression of the GFP reporter, but levels of Vpr targeted REAF and MUS81 protein remain unchanged (Figure 1b). To make these cell lines permissive to HIV-1 infection, they were transfected with a lentiviral CD4 expression plasmid and cell surface CD4 expression was confirmed by FACS analysis (Figure S1). Monolayers of HeLa-CD4 and HeLa-ΔTASOR-CD4 cells were challenged with serial dilutions of HIV-1 89.6^WT^ and infectivity was determined by quantifying foci of infected cells with intracellular staining of viral p24 protein (Figure 1c). HeLa-ΔTASOR-CD4 were significantly (p<0.0001) more susceptible to infection than HeLa-CD4 (Figure 1c). Interestingly, we did not detect enhanced susceptibility in the absence of HuSH complex member Periphilin (HeLa-ΔPPHLN1-CD4). Implying the existence of a TASOR mediated restriction to HIV-1 replication independent of HuSH silencing activity which requires Periphilin.

Given the ability of HIV-1 Vpr to downmodulate TASOR (Figure 1a) and the presence of a TASOR mediated restriction in HeLa-CD4 cells (Figure 1c), we hypothesised that removal of TASOR would be more advantageous to a virus lacking Vpr (HIV-1 89.6^Δ*vpr*^) than to wild-type virus (HIV-1 89.6^WT^).

To test this hypothesis, the infectivity of HIV-1 89.6^WT^ and HIV-1 89.6^Δ*vpr*^ on HeLa-CD4 and HeLa-ΔTASOR-CD4 was determined (Figure 1d). We also challenged cells with mutant viruses containing amino acid substitutions within Vpr - HIV-1 89.6^Q65R^ and HIV-1 89.6^F34I^. The F34I mutant Vpr is partially defective at localising to the nuclear membrane and at interacting with the nuclear transport proteins importin-α and nucleoporins while Q65R abrogates Vpr-DCAF1 binding and impairs Vpr docking at the nuclear envelope ^45–48^. Both mutations increase the susceptibility of the virus to TASOR (also previously shown for REAF ^6^) which is predominantly nuclear localised (Figure S2). All viruses could replicate more efficiently in the absence of TASOR (HeLa-ΔTASOR-CD4 compared to HeLa-CD4), but the fold increase in virus replication was significantly greater for HIV-1 89.6^Δ*vpr*^ (p <0.0001), HIV-1 89.6^Q65R^ (p=0.0039) and HIV-1 89.6^F34I^ (p=0.0039) when compared to HIV-1 89.6^WT^. To further confirm these results we generated cell lines with stable expression of shRNA specific for TASOR (HeLa-shTASOR-CD4) or REAF (HeLa-shREAF-CD4). TASOR restriction in these cells was tested by titration of primary isolate HIV-1 2044 ^49^. Infectivity of HIV-1 2044 was significantly greater in both HeLa-shTASOR-CD4 (p=0.0045) and HeLa-shREAF-CD4 (p<0.0001) where 10-30-fold more FFU/ml were detected compared to control cells (Figure 1e). Also, the infectivity of HIV-1 89.6^Δ*vpr*^ and HIV-1 89.6^WT^ in these cells was compared with the infectivity of a virus lacking another accessory gene *vpu* (HIV-1 89.6^Δ*vpu*^). Depletion of TASOR by RNAi (or REAF as a positive control) resulted in enhanced infectivity for all viruses tested, but the enhancement was significantly greater for HIV-1 89.6^Δ*vpr*^ compared to HIV-1 89.6^WT^ (Figure 1f). Conversely, there was no enhancement in the infectivity of HIV-1 89.6^Δ*vpu*^.

Given the importance of Vpr for infection of macrophages and their importance in the pathology of HIV-1 infection, we investigated TASOR restriction in primary monocyte derived macrophages (MDMs) ^28,30^. Treatment of MDMs with VLPs containing HIV-1 Vpr resulted in a loss of TASOR protein (Figure 1g). Analysis of subcellular TASOR protein levels by imaging flow cytometry further revealed that the loss of TASOR was predominantly from the nuclear region of the cell as Vpr-VLP treated MDMs have a significantly lower TASOR/DAPI Similarity Score (p=0.0286), a measure of chromatin associated TASOR compared to TASOR in the cell overall (Figure 1h). Representative images of TASOR in MDMs with DAPI stained nuclei treated with either empty VLPs (left) or Vpr-VLPs (right) are shown in Figure 1i where an absence of TASOR can be seen in the nucleus (DAPI) of only the Vpr-VLP treated MDMs. The subsequent infectivity of these MDMs was assessed by challenge with either HIV-1 89.6^WT^ or HIV-1 89.6^Δ*vpr*^, virus production was compared by ELISA of HIV-1 p24 protein in cell culture supernatants (Figure 1j). There was an increase in production of both HIV-1 89.6^WT^ and HIV-1 89.6^Δ*vpr*^ from MDMs after prior Vpr-VLP treatment. However, the increase was significantly greater (p=0.0079) for HIV-1 89.6^Δ*vpr*^ compared to HIV-1 89.6^WT^ suggesting Vpr mediated TASOR depletion is more advantageous to a virus lacking Vpr of its own.

### HIV-1 Vpr overcomes early TASOR mediated restriction to reverse transcription

Vpr and Vpx are specifically incorporated into the virus particle via the p6 domain of the Gag precursor and released into target cells rapidly after entry suggesting they have a role prior to integration of the viral genome ^24,50^. Vpr locates to the nucleus within 45 minutes of viral challenge ^51^. The kinetics of REAF protein depletion early after infection have been previously studied, there is a Vpr mediated loss seen as soon as 30 minutes post challenge with transient recovery possibly due to the limited amount of Vpr in the virus particle and high protein turnover ^5,6^. Here we show that Vpr mediated TASOR protein down-modulation also occurs rapidly after infection (Figure 2a). From just 2 hours following challenge with HIV-1 89.6^WT^, a loss of TASOR protein is observed which is sustained throughout the time course up to 24 hours. Conversely, TASOR protein levels do not change when the infecting virus lacks Vpr (HIV-1 89.6^Δ*vpr*^). Analysis of other Vpr-depleted cellular proteins MUS81 and HLTF over time post viral challenge shows that, like REAF and TASOR, the loss of protein also occurs within one hour after HIV-1 89.6^WT^ infection (Figures 2b, c). In contrast to REAF, HLTF and MUS81 levels remain low.

**Figure 2:**
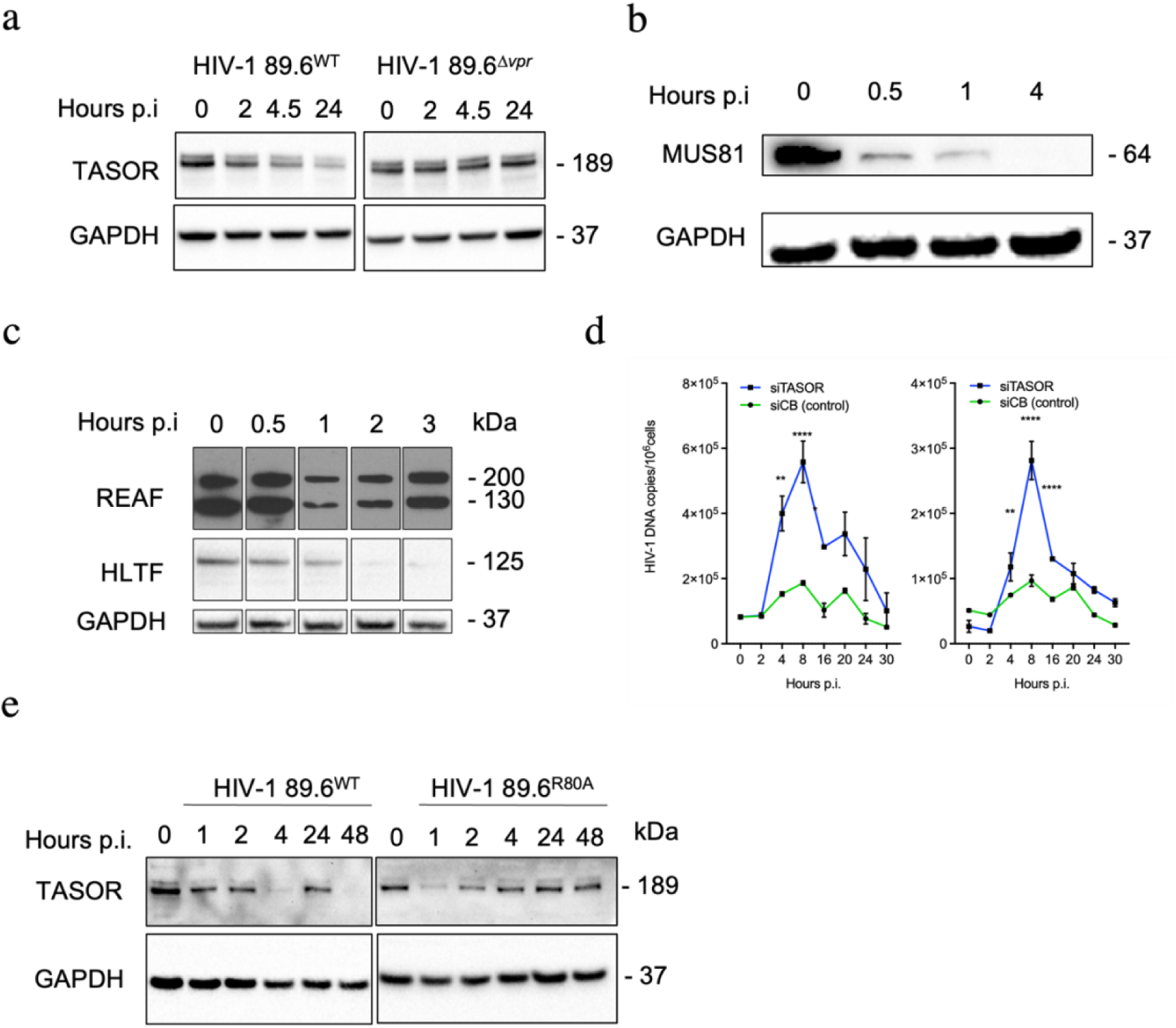
HIV-1 Vpr overcomes early TASOR mediated restriction to reverse transcription. **a**, Western blotting of TASOR in HeLa-CD4 cells over time (0-24 hours) after challenge with HIV-1 89.6^WT^ or HIV-1 89.6^Δ*vpr*^. **b**, Western blotting of MUS81 protein levels in THP-1 cells over the first 4 hours post challenge with HIV-1 89.6^WT^. **c**, Western blotting of REAF and HLTF protein levels in THP-1 cells over the first 3 hours post challenge with HIV-1 89.6^WT^ (full blot +/- exogenous dN is in Figure S3). **d**, qPCR HIV-1 RT products – strong stop (left) and late (right) – 0–30 hr post infection (p.i.). Cells were treated with siRNA targeting TASOR (siTASOR) or a control gene Cyclophilin B (siCB). HIV-1 DNA copies measured by qPCR are normalised to genomic GAPDH and presented per 10^6^ cells. Error bars represent standard deviations of means of duplicates. **e**, Western blotting of TASOR in PMA differentiated THP-1 cells over time (0-48 hours) after challenge with HIV-1 89.6^WT^ or HIV-1 89.6^R80A^. p.i., post infection.

We previously showed that REAF restricts the replication of HIV-1 resulting in the production of fewer reverse transcripts in the infected cell and that REAF mediated restriction of viral replication is overcome by HIV-1 Vpr by promoting REAF degradation ^5,6^. Similarly, we now show using qPCR that in the absence of TASOR there are significantly elevated levels of both early (strong-stop) and late products of reverse transcription (Figure 2d). Between 4h and 8h (strong stop, left) and between 4h and 16h (late, right) copy numbers are greater in siTASOR treated cells compared to cells transfected with siRNA targeting a control gene (Cyclophilin B) (Figure 2d).

The ability of Vpr to induce arrest at G2/M in a cycling population of cells, but not its ability to bind DCAF1, is disrupted by the introduction of an alanine in place of arginine at position 80 (R80A) ^52,53^. We challenged THP-1 cells with either HIV-1 89.6^WT^ or HIV-1 89.6^R80A^ and harvested cells over time for analysis of TASOR protein levels by Western blotting. TASOR protein levels are almost undetectable by 4 hours post challenge with HIV-1 89.6^WT^ (Figure 2e, left). We observe a transient recovery of TASOR detected at 24 hours, prior to further downmodulation after integration. Interestingly, while an early loss of TASOR in the initial phase was detected after infection with HIV-1 89.6^R80A^ levels are rapidly recovered and no further loss is detected (Figure 2e, right).

### TASOR depletion causes cell cycle arrest and arrested cells are more susceptible to infection

Vpr induces cell cycle arrest at G2/M after infection and depletion of Vpr targets including REAF and others such as CCDC137 results in cell cycle perturbation ^6,35,37,54,55^. The extent of cell cycle arrest induced by Vpr may only be partially explained by downmodulation of REAF and it is hypothesised by others that more than one protein may be required to produce the strong Vpr-induced G2/M arrest phenotype reported ^6,35^. To test whether downmodulation of TASOR contributes to the cell cycle perturbation observed after infection with a virus that has Vpr, DAPI stained HeLa and HeLa-ΔTASOR were analysed by imaging flow cytometry for DNA content. The cell cycle profiles show an accumulation of cells in the G2/M phase of the cell cycle with double DNA content in HeLa-ΔTASOR compared to HeLa (Figure 3a). Complete cell cycle arrest was not expected in these CRISPR-Cas9 knock out cells as they proliferate in culture despite TASOR depletion.

**Figure 3:**
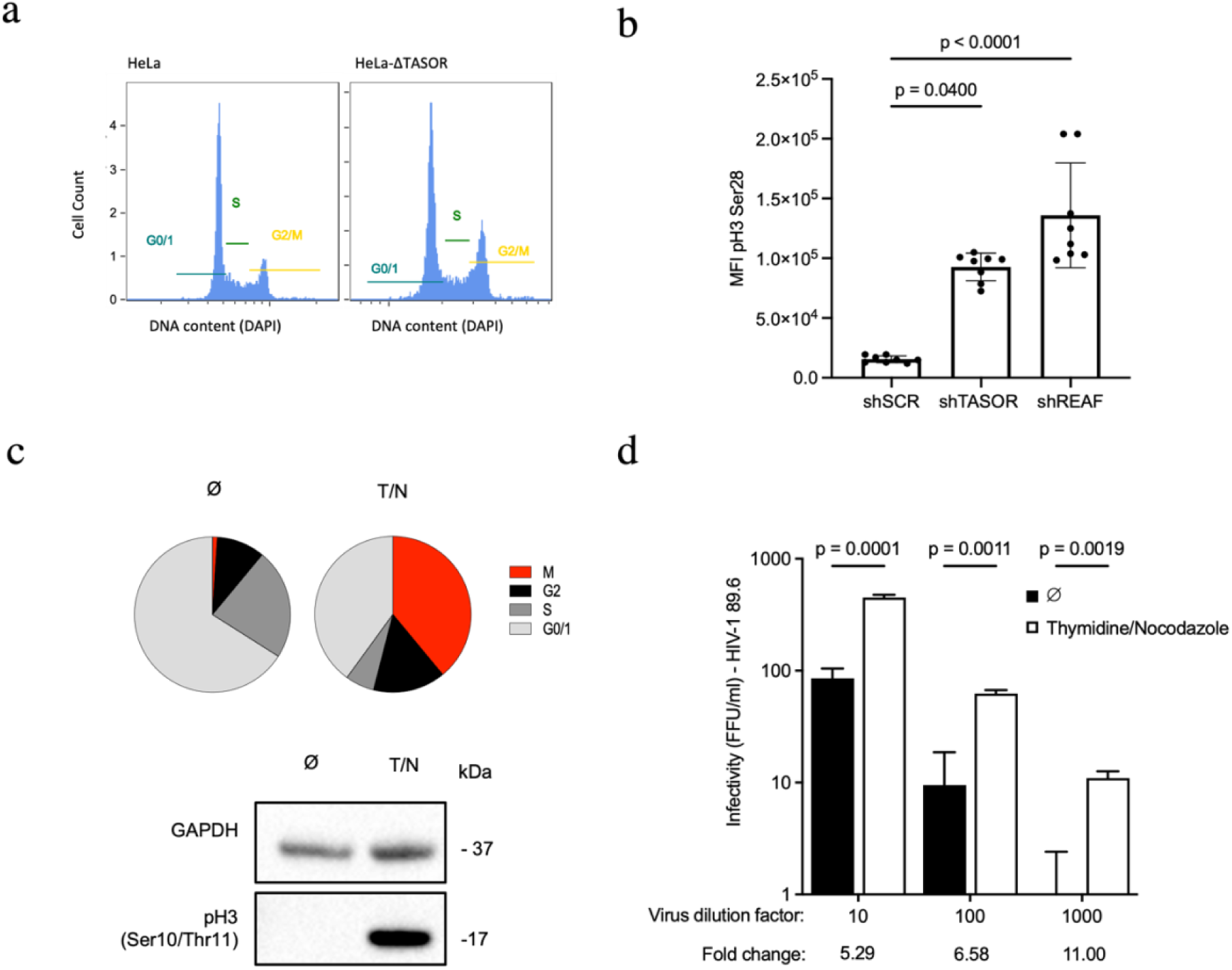
TASOR depletion causes cell cycle arrest and arrested cells are more susceptible to infection. **a**, Cell cycle analysis of DAPI stained HeLa (left) and HeLa-ΔTASOR (right) by imaging flow cytometry. **b**, Imaging flow cytometry of phosphorylated histone H3 (pH3 Ser28) in HeLa-shSCR, HeLa-shTASOR and HeLa-shREAF. Error bars represent the standard deviations of means of replicates. Kruskal-Wallis multiple comparison and ANOVA with Dunn’s correction. **c**, Proportions of untreated HeLa-CD4 (Ø) and thymidine/nocodazole (T/N) treated HeLa-CD4 cells in different phases of the cell cycle (G0/1, S/ G2, M) determined by flow cytometry of cells with DAPI staining for DNA quantification and staining of phosphorylated histone H3 Ser28 to identify cells in the mitotic phase (M). Western blotting of phosphorylated histone H3 Ser10/Thr11 in the same cells. **d**, Infectivity (FFU/ml) of HIV-1 89.6^WT^ in untreated HeLa-CD4 (Ø) and thymidine/nocodazole (mitotic cell enriched) HeLa-CD4 cells at three different virus inputs (dilution factor indicated). Fold changes in susceptibility to infection are shown. Error bars represent the standard deviations from the means of technical replicates. Unpaired t-tests.

To further probe the relationship between loss of TASOR and Vpr mediated cell cycle arrest in the G2/M phase, we measured levels of mitotic cell marker phosphorylated histone H3 (Ser28) by imaging flow cytometry after TASOR knock down (HeLa-shTASOR-CD4). Mean fluorescence intensity (MFI) of pH3 Ser28 was significantly greater in HeLa-shTASOR-CD4 (p = 0.04) and in HeLa-shREAF-CD4 (p < 0.0001) compared to HeLa-shSCR-CD4 (Figure 3b).

Despite the well-established association between HIV-1 Vpr and cell cycle arrest, the advantage to the virus is not fully understood ^26,54,55^. We hypothesised, given the partial cell cycle arrest observed after TASOR knock down, that mitotic cells may be more susceptible to infection. To test this, we challenged a mitotic enriched population of cells with replication competent HIV-1 89.6^WT^. A mitotic cell enriched population of HeLa-CD4 was achieved using a thymidine-nocodazole synchronisation protocol and mitotic enrichment was quantified using flow cytometry and Western blotting. In the treated population, 54% (red and black) were in the G2/M phase and 39% (red) were in mitosis (with high levels of pH3 Ser28) (Figure 3c). In the untreated population, only 11% (red and black) were in the G2/M phases and only 1% (red) were in mitosis (Figure 3c). Western blotting also confirmed mitotic enrichment with the synchronised thymidine/nocodazole cells also demonstrating high levels of mitotic marker pH3 Ser10/Thr11 (Figure 3c). HIV-1 89.6^WT^ was titrated on both treated and untreated cells and staining for foci of infection revealed that the mitotic enriched population was approximately 5-11 fold more susceptible to infection (Figure 3d). At each virus dilution the difference in infectivity between mitotic-arrested and cycling was significant.

## Discussion

In this study we have demonstrated a novel restrictive activity of TASOR which results in impaired reverse transcription of viral RNA early in infection. In cells silenced for TASOR expression, an increase in RT products is seen within 4 hours following viral challenge. We show that TASOR restriction is potent in primary macrophages, active against a primary HIV-1 isolate and mitigated by Vpr. While cells lacking TASOR were more susceptible to infection, those lacking HuSH component PPHLN1 were not - suggesting the existence of a restriction involving TASOR that is independent of HuSH mediated repression of HIV-1 post integration. Thus, like other restriction factors such as SAMHD1, TASOR challenges the virus at multiple stages of the replication cycle.

Counteracting this restriction, HIV-1 Vpr downmodulates TASOR protein within 1-2 hours of infection, coinciding with the initiation of reverse transcription as well as 48 hours later when *de novo* synthesis has occurred. Others have reported a segregation of Vpr packaged in progeny virions and those of *de novo* expressed Vpr later in the virus life cycle ^56^. The detection of a TASOR mediated restriction provides an explanation for the early loss of TASOR protein induced by Vpr. It also offers an explanation as to why Vpr is actively incorporated into the virus particle with high copy numbers as it is needed prior to production of *de novo* Vpr ^57^.

Macrophages are critical players in HIV-1 infection and persistence ^58–60^. They are also the dominant source of viremia after the reduction in the CD4+ T cell population and HIV persists in myeloid cells with high frequency ^28,61^. We show that TASOR is restrictive to replication competent virus in terminally differentiated macrophages and that HIV-1 Vpr alone, delivered by VLP, is sufficient to down-modulate TASOR in these cells. In further agreement with the proposition that HIV-1 Vpr alleviates TASOR restriction in macrophages, pre-treatment of MDMs with VLPs containing Vpr prior to infection is more advantageous to a replicating virus lacking Vpr (HIV-1 89.6^Δ*vpr*^). This is the first report of HIV-1 Vpr mediated counter restriction involving TASOR and these findings are further evidence of the multifaceted functionality of HIV-1 Vpr and the importance of this accessory protein for HIV-1 replication.

TASOR is a predominantly nuclear protein and Vpr rapidly localises to the nucleus in the same timeframe that we see loss of TASOR protein after infection ^51^. Given the similarity between the TASOR phenotype described here and that of REAF described previously we propose that both REAF and TASOR act together. Recently, it was shown by proximity dependent labelling (BioID) that TASOR associates with REAF and 9 other chromatin localised proteins ^8^. The increased susceptibility of HIV-1 89.6^F34I^ to TASOR restriction confirms that Vpr localisation and its association with nuclear transport machinery is important for counter restriction of TASOR. The nucleus is now arguably the primary site of reverse transcription and several studies have determined that that reverse transcription is completed within the intact capsid core which uncoats less than 1.5μm from the site of integration ^62–64^. HIV-1 Vpr Q65R mutant is also more susceptible to TASOR mediated restriction, confirming that down-modulation is achieved utilising the cellular proteasome and DCAF1, a substrate recognition component of the CRL4-DCAF1 E3 ligase complex^48^. These experiments support our proposition that virion incorporated Vpr carried in the virus particle rapidly locates to the nucleus and degrades TASOR to alleviate a block to reverse transcription.

In infection time course experiments, we found that *de novo* produced Vpr can also downmodulate TASOR but the R80A mutant cannot. The c-terminal domain HIV-1 Vpr mutant R80A retains the ability to bind to DCAF1 but can no longer induce G2/M arrest ^52^. This result indicates that Vpr mediated mitigation of TASOR early in infection may be mechanistically distinct from TASOR activity post integration and from the G2/M arrest phenotype for which Vpr is known. Furthermore, the inability of the R80A mutant to downmodulate TASOR later in infection genetically separates the early and late effects of Vpr/TASOR. While we cannot definitively link Vpr mediated cell cycle arrest and TASOR degradation, our data suggests that they may be associated. This is supported by analysis of TASOR depleted cells which reveals there is an accumulation in G2/M. Cycling cells lacking TASOR have a higher proportion of the population in the G2/M phase with double DNA content and expressing high levels of the mitotic marker phosphorylated histone H3 (Ser28). We also show that cells arrested in mitosis are more susceptible to infection with HIV-1 than a cycling population. Complete cell cycle arrest is not observed, and it is likely that degradation of multiple factors is required.

Taken together, the results presented here support the current model for Vpr activity, which is that it is actively incorporated into the virus particle so that it can overcome an early block to reverse transcription and induce the degradation of proteins involved in restriction of HIV-1. TASOR is a crucial component of a Vpr-targeted restriction system that is active against HIV-1 both early and late in infection. Disrupting the ability of Vpr to overcome TASOR may be an important therapeutic strategy.

## Methods

### Cell Lines and Culture Reagents

HEK-293T, Jurkat, THP-1 and HeLa cell lines were cultured in Dulbecco’s Modified Eagle Medium (DMEM) and C8166 were cultured in Roswell Park Memorial Institute Medium (RPMI-1640) supplemented with 5-10% heat-inactivated foetal bovine serum (FBS), 0.1% 200mM L-glutamine, sodium pyruvate and appropriate antibiotics. Cultures were periodically cultured in media supplemented with penicillin/streptomycin at 1U/ml and 0.1 mg/ml respectively. All cell lines were maintained and cultured at 37°C in 5% CO_2_. THP-1 cells were differentiated into a macrophage-like state by treatment in culture with phorbol 12-myristate 13-acetate (PMA) at a concentration of 62ng/ml for 72 hours and then PMA-free DMEM supplemented with 10% FBS and Penicillin/streptomycin for a further 48 hours to allow further differentiation and recovery prior to infection or analysis.

CRISPR-Cas9 HeLa cell lines with a repressed GFP reporter, including those deficient for TASOR (HeLa-ΔTASOR) or Periphilin (HeLa-ΔPPHN1), were a gift from Prof. Paul Lehner (University of Cambridge, United Kingdom) ^7^. To make these cell lines permissible to infection, they were transfected with CD4 expression plasmid pCMS28 ENX to generate puromycin resistant HeLa-CD4, HeLa-ΔTASOR CD4 and HeLa-ΔPPHN1-CD4. Puromycin resistant colonies were selected and maintained in culture with puromycin. The concentration of puromycin required for selection and maintenance was determined in dose response experiments. Surviving cells at each concentration were stained using crystal violet solution (0.1% w/v) and 570nm absorbance of the stain, resuspended using 1% w/v sodium dodecyl sulphate (SDS), was determined using a CLARIOStar Plate Reader (BMG Labtech). The typical concentration required for complete kill was between 5 μg/ml and 15μg/ml. Clonal colonies of cells resistant to concentrations of puromycin greater than that required for complete kill of the non-transfected cells were selected and subjected limiting-dilution clone selection. After selection, cell lines were cultured under continuous puromycin presence. An example puromycin ‘kill-curve’ on non-transfected cells is shown in Figure S1a.

The preparation of HeLa-CD4 cell lines with stable expression of short hairpin RNA (shRNA) has previously been described ^65^. The pSUPER RNA interference system (pSUPER.retro.puro; Oligoengine) was used for TASOR and REAF depletion in HeLa-CD4 (HeLa-CD4-shTASOR/shREAF). The lentiviral shRNA expression vector was either transfected directly into cells or delivered by preparation of retroviral particles. Briefly, HEK-293T cells were co-transfected with shRNA expression plasmid, HIV-1 gag-pol expression vector (p8.91) and the vesicular stomatitis virus G protein (VSVG) expression vector pMDG VSV-G. Viral vector containing supernatant was harvested after 48h, clarified by centrifugation at 500xg and passed through a 0.45μm pore size filter. HeLa-CD4 were transduced by culturing with retroviral supernatants for 72 hours, after which resistant colonies were selected and maintained with puromycin. The concentration of puromycin required for selection and maintenance was determined in dose response experiments (Figure S1a).

### Transfections, Virus and VLP Production

The infectious molecular clones for HIV-1 89.6^WT^ were obtained from the Centre for AIDS Research (NIBSC, UK). The HIV-1 89.6^Δvpr^ plasmid construct was generated from the HIV-1 89.6^WT^ molecular clone, using overlap extension PCR. HIV-1 89.6 *vpr* mutants F341, R80A and Q65R were made by site directed mutagenesis (Agilent) of the HIV-1 89.6^WT^ plasmid. Clones were confirmed by plasmid sequencing (Source BioScience). The virus-like particle (VLP) system pMDLchp6, pcDNA3.1-HIV-1-Vpr and pcDNA3.1-SIVmac239-Vpx were a gift from N. Landau ^30,66,67^. Infectious molecular clones were amplified where appropriate by addition of C8166 prior to harvest. For virus/VLP production, HEK-293T cells were plated at 2×10^4^/cm^2^ in 60cm^2^ cell culture dishes 48 hours prior to transfection. Infectious full-length, chimeric HIV clones and VLPs were prepared by transfection of HEK-293T cells using Lipofectamine 3000 (Invitrogen) or Fugene 4K (Promega). The HIV-1 primary isolate 2044 (described previously ^68^) was isolated from the peripheral blood of a Clade B HIV-infected patient and minimally passaged in in PBMCs from healthy donors. Fresh PBMCs were plated in 10cm^2^ cell culture dishes and inoculated with 5×10^4^ FFU in 5ml DMEM supplemented with 10% FBS and penicillin/streptomycin. Viral supernatants were harvested after 48 and 72 hours and cleared of cell debris by centrifugation at 500 x g for 5 minutes. Virus stocks were stored at -196°C.

### Virus Titrations

For all infectivity experiments conducted comparing HIV-1 and mutants on cell lines with and without REAF/TASOR, viral input was normalised across variants by first calculating the infectivity of each viral stock on wild-type cells. Multiplicity of infection (MOI) in each instance is stated and was equivalent when comparing viruses or comparing the infectivity of one virus on different cell lines/types. Where the titre of these viruses is expressed as FFU/ml, titre was calculated in HeLa-CD4 seeded at 1.5×10^4^ cells/well in 48-well plates 24 hours prior to virus titration. Cell monolayers were challenged with serial 1 in 10 dilutions of virus stock and the titre was assessed after 48 hours by in situ intracellular staining of HIV-1 p24 to identify discrete foci of viral replication. For in situ intracellular staining, cell culture supernatants were aspirated from wells and the cells were washed in PBS. Cells were then fixed in ice-cold methanol:acetone (50:50) and p24 antigen was stained using mAb to p24 diluted 1 in 40 (1:1 mix of EVA 365 and 366 from the MRC AIDS Reagent Program, UK) in PBS 1% FBS for 1h at 37°C. After incubation with the primary antibody, cells were washed a further 3 times in PBS and incubated with goat anti-mouse IgG β-galactosidase-conjugated antibody (Southern Biotech) diluted 1 in 400 in PBS 1% FBS for 1h at 37°C. After 3 further PBS washes, 300 μL of 0.5mg/ml 5-bromo-4-chloro-3-indolyl ß-D-galactopyranoside chromogenic substrate (X-gal) in PBS containing 3 mM potassium ferricyanide, 3 mM potassium ferrocyanide and 1 mM magnesium chloride was added to each well. Infected cells incubated at 37°C and stained blue within 1 to 4 hours after addition of substrate. Foci of infection were counted to determine the virus titre defined as focus forming units per ml (FFU/ml).

### Infections and VLP Treatment

For infection time course experiments, 500 μL of 1×10^5^ FFU/ml virus was added per well to cells cultured in 6-well trays. MOI was approximately 0.25. Cells were harvested with trypsin over time for analysis by Western blotting. For VLP experiments and experiments with MDMs where viral production was analysed, cells were challenged with equal viral/VLP inputs of 50ng p24, determined by enzyme-linked immunosorbent assay (ELISA), in 6-well plates with 2×10^6^ cells per well. Total volume was 750μl. Supernatants were harvested and p24 concentration was also assessed by ELISA.

For time course infection experiments analysed by qPCR, HeLa-CD4 cells were seeded at 2.5 × 10^4^ cells/well in 24-well plates. siRNA transfection (30nM) was performed using HiPerfect (QIAGEN) according to the manufacturer’s instructions. 72 hr after siRNA transfection, cells were challenged with virus (MOI 0.2). Infection was assessed up to 48 hr by qPCR analysis.

### HIV-1 p24 Antigen ELISA

ELISA plates were pre-coated with 5μg/ml sheep anti-HIV-1 p24 antibody (Aalto Bio Reagents) at 4°C overnight. Viral supernatants treated with 1% Empigen® BB at 56°C for 30 minutes, were then plated at a 1:10 dilution in Tris-buffered saline (TBS) on anti-p24-coated plates and incubated for 3 hours at room temperature. Alkaline phosphatase-conjugated mouse anti-HIV-1 p24 monoclonal antibody (Aalto Bio Reagents) diluted in TBS 20% sheep serum, 0.05% v/v Tween-20 was then added and incubated for 1 hour at room temperature. Plates were washed 4 times with 0.01% v/v Tween-20 in PBS and twice with ELISA Light washing buffer (ThermoFisher). CSPD substrate with Sapphire II enhancer (ThermoFisher) was added and incubated for 30 minutes at room temperature before chemiluminiscence was detected using a CLARIOStar Plate Reader (BMG Labtech). p24 antigen titres obtained using this method were used to normalise inputs of VLPs used in challenge experiments.

### Western Blotting

Adherent cells were harvested with trypsin and lysed in 30-50 μL of radioimmunoprecipitation assay (RIPA) buffer supplemented with NaF (5μM), Na_2_VO_3_ (5μM), β-glycerophosphate (5μM) and 1x Protease Inhibitor Cocktail (Cytoskeleton). The protein concentration of each sample was determined using BCA Protein Assay Kit (Pierce). 12.5μg of total protein was separated by SDS-PAGE (4-12% Bis-Tris Gels, Invitrogen), at 120V for 1 hour 45 minutes in MOPS SDS Running Buffer (Invitrogen). Separated proteins were transferred onto nitrocellulose membrane (0.45μm pore size, GE Healthcare) at 45V for 2 hours, in ice-cold 20% (v/v) Methanol NuPAGE™ Transfer Buffer (ThermoFisher). After transfer, membranes were stained for total protein using Ponceau S staining solution (0.1% (w/v) Ponceau in 5% (v/v) acetic acid), washed 3 times for 5 minutes on an orbital shaker in dH_2_O and imaged using ChemiDoc Gel Imaging System. Membranes were blocked for 1 hour at room temperature in 5% (w/v) non-fat milk powder in TBS-T buffer or 2.5% (w/v) bovine serum albumin (BSA). Specific proteins were detected with primary antibodies by incubation with membranes overnight at 4°C and with secondary antibodies for 1 hour at room temperature. All antibodies were diluted in blocking buffer. Proteins were visualized using ECL Prime Western Blotting Detection Reagent (GE Healthcare) and imaged using either ChemiDoc Gel Imaging System (Bio-Rad) or exposed to CL-XPosure films (ThermoScientific) and developed. Detection of GAPDH and membrane staining with Ponceau S solution was used to confirm equal loading of total protein and for normalisation in densitometry analysis. Densitometry was performed using either ImageJ for Mac or Image Lab software version 6.0 for PC.

### Antibodies

For imaging flow cytometry RbpAb-TASOR/FAM208A (Atlas Antibodies) was detected using goat anti-rabbit IgG conjugated with Alexa Fluor 647 (Invitrogen). FITC-labelled anti-phospho-histone H3 (Ser28) Alexa 488 was used (BD Bioscience) for imaging flow cytometry. APC-mouse anti-human CD4 was used for flow cytometry (clone RPA-T4, BioLegend). MsmAb-MUS81 (Abcam) was detected by anti-mouse IgG antibody conjugated to HRP (GE Healthcare) for Western blotting. RbpAb-REAF/RPRD2 (Eurogentec), RbpAb-TASOR/FAM208A (Atlas Antibodies), RbpAb-GAPDH (Abcam), RbpAb-HLTF (Abcam) and RbmAb-phospho-histone H3 Ser10/Thr11 (Abcam) were detected with donkey anti-rabbit IgG conjugated to HRP (GE Healthcare) for Western blotting. For in situ intracellular staining of HIV-1 mAb to p24 diluted 1 in 40 (1:1 mix of EVA 365 and 366 from the MRC AIDS Reagent Program, UK) was used and detected using goat anti-mouse IgG β-galactosidase-conjugated antibody (Southern Biotech) diluted 1 in 400.

### Isolation of PBMCs from Peripheral Blood

PBMCs were isolated from leukocyte cones containing peripheral blood (NHS Blood Transfusion service, St. George’s Hospital, London) by density gradient centrifugation with Lymphoprep™ (STEMCELL™ Technologies) or Ficoll® Paque Plus (MERCK) density gradient medium. Peripheral CD14+ monocytes were isolated from PBMCs using the human CD14+ magnetic beads (Miltenyi Biotech) according to the manufacturer’s instructions. To obtain monocyte derived macrophages (MDMs), monocytes were treated with 100ng/ml macrophage colony stimulating factor (M-CSF, Peprotech) for 7 days, with medium replenished on day 4. Magnetic beads were removed prior to intracellular staining and imaging flow cytometry.

### qPCR of HIV-1 Reverse Transcripts

DNA was extracted from cells at various time points after infection with a QIAamp DNA Blood Mini Kit (QIAGEN). The isolated DNA was subjected to real-time quantitative PCR (qPCR) to determine the number of early (negative strand strong stop, -sss) and late (gag) transcripts present, normalised for cell number by genomic GAPDH. This methodology has been previously described ^69^.

### Flow Cytometry and Imaging Flow Cytometry

Cells analysed for CD4 expression by flow cytometry were incubated with Fc block, TruStain FcX (1:50, BioLegend) in FACS buffer (1mM EDTA, 2% FCS in PBS) for 15 minutes at room temperature, then incubated with APC-mouse anti-human CD4 (0.2 microgram/ml, clone RPA-T4, BioLegend) for 20 minutes at room temperature. Cells were then washed with FACS buffer and fixed with 2% PFA/PBS for 30 minutes at room temperature, before analysis with flow cytometer LSR II (BD).

Cells analysed by imaging flow cytometry were fixed in FIX&PERM® Solution A (Nordic MUbio) for 30 minutes and permeabilised with 0.2% Triton™-X 100/PBS at room temperature. MDMs were blocked with human serum (1%) for 1 hour prior to staining. The staining buffer used was PBS 0.1% Triton™-X 100, 0.5% FBS. Nuclei were counterstained with DAPI (1μg/ml) for two hours prior to analysis. Fixed and stained cells were stored at 4°C in PBS with sodium azide (0.1%). Imaging flow cytometry was performed using the Amnis ImageStream®x Mark II Flow Cytometer (Merck) and INSPIRE® software (Amnis). A minimum of 5000 events were collected per sample. IDEAS® software (Amnis) was used for analysis and to determine subcellular protein levels and similarity score. Single cells were distinguished from aggregates and selected for inclusion in the analysis by generating a scatter plot of brightfield area vs brightfield aspect ratio. Aspect ratio values relate to the shape of the cells and is the ratio of cellular minor axis (width) to major axis (height). Of the single cells, only those that were in-focus were gated by plotting a histogram of gradient root mean squared for image sharpness. Cells stained for target protein and DAPI were further gated from the in focus single cells.

Subcellular TASOR protein levels were quantified within the nucleus and cytoplasm using DAPI staining to define the nuclear region and brightfield images, from which the nuclear region was subtracted to define the cytoplasmic region. The similarity score feature is derived from Pearson’s correlation coefficient (pixel-by-pixel correlation of the TASOR and nuclear dye image pair within the nuclear morphology mask dilated by one pixel) and relates to the degree of co-localisation.

### Synchronisation and Infection of Mitotic Cells

To synchronise cells in the mitotic phase of the cell cycle, an initial S phase block with thymidine (4mM) was induced for 24 hours followed by a PBS wash and treatment with nocodazole (100ng/ml) for a further 16 hours. Cells in suspension and those detached easily with a manual “shake off” enriched the population of cells in the mitotic phase ^70^. Cells were allowed to cycle for two hours in culture before infection with HIV-1 89.6^WT^ in suspension in 48-well plates. Virus was titrated against 5×10^4^ cells in a volume of 400μl. Infectivity was determined after 48 hours by intracellular staining of p24 to detect and quantify foci of infection.

### Cell Cycle Analysis

Cell cycle phase distribution of cells was determined by analysis of DNA content by imaging flow cytometry. Cells were fixed in FIX&PERM® Solution A (Nordic MUbio) and stained with DAPI (1μg/ml) before analysis. Mitotic cells were detected by imaging flow cytometry using the antibody FITC-labelled anti-phospho-histone H3 (Ser28) Alexa 488 to detect phosphorylated histone H3, a marker of mitotic cells. Cell lysates were also assessed by Western blotting using the anti-phospho-histone H3 (Ser10/Thr11) antibody as an additional marker of mitotic cells.

### Statistics and Reproducibility

Statistical significance was calculated using statistical tests stated in figure legends, in all cases, a p value of < 0.05 was considered statistically significant. Error bars represent standard deviations from the means of biological replicates. Prism software version 9 and Excel were used for analysis and illustration of graphs.

## Acknowledgements and Funding

The monoclonal antibodies to HIV-1 p24 (EVA365 and EVA366) were provided by the EU Program EVA Centre for AIDS Reagents, NIBSC, United Kingdom (AVIP contract number LSHP-CT-2004-503487). We thank Maximilien Biguet for discussions and helpful suggestions. We thank Prof. N. Landau for the kind gift of VPL constructs and Prof. P. Lehner for the kind gift of HeLa cell lines. J.M.G and A.M.K received funding from the Rosetrees Trust (M665 and CF1\10003) and Barts Charity (G-002017). This research was also supported by the BCI Flow Cytometry Facility at Barts Cancer Institute (CRUK Flow Cytometry Core Service Grant C16420/A18066). The Wellcome Trust (101604/Z/13/Z) funded the purchase of an Amnis ImageStream imaging flow cytometer. The funders had no role in the study design, data collection, data analysis, data interpretation, decision to publish or preparation of the manuscript.

## Data Availability

The datasets generated during and/or analysed during the current study are available from the corresponding author on reasonable request.

## Ethical Approval

Leucocyte cones, from which peripheral blood mononuclear cells (PBMCs) were isolated and monocyte-derived macrophages (MDMs) cultured, were obtained from the NHS Blood Transfusion service at St. George’s Hospital, London. Donors were anonymised and thus patient consent was not required. The local ethical approval reference number is 06/Q0603/59.

## Author Contributions

Á.M and J.M.G conceived the project and obtained funding. J.M.G., K.M., C.P., W-Y. J. L. and Á.M designed experiments. J.M.G., K.M., C.P. and W-Y. J. L. performed experiments. J.M.G., K.M., C.P., W-Y. J. L. and Á.M analysed and interpreted the data. J.M.G prepared the figures. J.M.G and Á.M. wrote the manuscript. Á.M, K.M., C.P. and W-Y. J. L. critically revised the manuscript. All authors reviewed and approved the manuscript and figures.

## References

1. Colomer-Lluch, M., Gollahon, L. S. & Serra-Moreno, R. Anti-HIV Factors: Targeting Each Step of HIV’s Replication Cycle. Curr HIV Res 14, 175–182 (2016).

2. Duggal, N. K. & Emerman, M. Evolutionary conflicts between viruses and restriction factors shape immunity. Nat. Rev. Immunol. 12, 687–695 (2012).

3. Simon, V., Bloch, N. & Landau, N. R. Intrinsic host restrictions to HIV-1 and mechanisms of viral escape. Nat Immunol 16, 546–553 (2015).

4. Liu, L. et al. A whole genome screen for HIV restriction factors. Retrovirology 8, 94 (2011).

5. Marno, K. M. et al. Novel restriction factor RNA-associated early-stage anti-viral factor (REAF) inhibits human and simian immunodeficiency viruses. Retrovirology 11, 3 (2014).

6. Gibbons, J. M. et al. HIV-1 Accessory Protein Vpr Interacts with REAF/RPRD2 To Mitigate Its Antiviral Activity. J Virol 94, (2020).

7. Tchasovnikarova, I. A. et al. GENE SILENCING. Epigenetic silencing by the HUSH complex mediates position-effect variegation in human cells. Science (80-.). 348, 1481–1485 (2015).

8. Douse, C. H. et al. TASOR is a pseudo-PARP that directs HUSH complex assembly and epigenetic transposon control. Nat. Commun. 11, 1–16 (2020).

9. Seczynska, M. & Lehner, P. J. The sound of silence: mechanisms and implications of HUSH complex function. Trends Genet. 39, 251–267 (2023).

10. Tchasovnikarova, I. A. et al. Hyperactivation of HUSH complex function by Charcot-Marie-Tooth disease mutation in MORC2. Nat. Genet. 49, 1035–1044 (2017).

11. Prigozhin, D. M. et al. Periphilin self-association underpins epigenetic silencing by the HUSH complex. Nucleic Acids Res. 48, 10313–10328 (2020).

12. Timms, R. T., Tchasovnikarova, I. A., Antrobus, R., Dougan, G. & Lehner, P. J. ATF7IP-Mediated Stabilization of the Histone Methyltransferase SETDB1 Is Essential for Heterochromatin Formation by the HUSH Complex. Cell Rep. 17, 653–659 (2016).

13. Zhu, Y., Wang, G. Z., Cingöz, O. & Goff, S. P. NP220 mediates silencing of unintegrated retroviral DNA. Nature 564, 278–282 (2018).

14. Matkovic, R. et al. TASOR epigenetic repressor cooperates with a CNOT1 RNA degradation pathway to repress HIV. Nat. Commun. 13, (2022).

15. Chougui, G. et al. HIV-2/SIV viral protein X counteracts HUSH repressor complex. Nat Microbiol 3, 891–897 (2018).

16. Yurkovetskiy, L. et al. Primate immunodeficiency virus proteins Vpx and Vpr counteract transcriptional repression of proviruses by the HUSH complex. Nat Microbiol 3, 1354–1361 (2018).

17. Colomer-Lluch, M., Ruiz, A., Moris, A. & Prado, J. G. Restriction Factors: From Intrinsic Viral Restriction to Shaping Cellular Immunity Against HIV-1. Frontiers in Immunology vol. 9 2876 (2018).

18. Hrecka, K. et al. Vpx relieves inhibition of HIV-1 infection of macrophages mediated by the SAMHD1 protein. Nature 474, 658–661 (2011).

19. Laguette, N. et al. SAMHD1 is the dendritic- and myeloid-cell-specific HIV-1 restriction factor counteracted by Vpx. Nature 474, 654–657 (2011).

20. Baldauf, H. M. et al. Vpx overcomes a SAMHD1-independent block to HIV reverse transcription that is specific to resting CD4 T cells. Proc. Natl. Acad. Sci. U. S. A. 114, 2729–2734 (2017).

21. Larrous, P. et al. Deciphering lentiviral Vpr/x determinants required for HUSH and SAMHD1 antagonism highlights the molecular plasticity of these evolutionary conflicts. bioRxiv 2024.03.07.583867 (2024) doi:10.1101/2024.03.07.583867.

22. Tristem, M., Purvis, A. & Quicke, D. L. J. Complex evolutionary history of primate lentiviral vpr genes. Virology 240, 232–237 (1998).

23. Selig, L. et al. Interaction with the p6 Domain of the Gag Precursor Mediates Incorporation into Virions of Vpr and Vpx Proteins from Primate Lentiviruses. J. Virol. 73, 592–600 (1999).

24. Bachand, F., Yao, X. J., Hrimech, M., Rougeau, N. & Cohen, E. A. Incorporation of Vpr into human immunodeficiency virus type 1 requires a direct interaction with the p6 domain of the p55 gag precursor. J Biol Chem 274, 9083–9091 (1999).

25. Paxton, W., Connor, R. I. & Landau, N. R. Incorporation of Vpr into human immunodeficiency virus type 1 virions: requirement for the p6 region of gag and mutational analysis. J. Virol. 67, 7229–7237 (1993).

26. Guenzel, C. A., Hérate, C. & Benichou, S. HIV-1 Vpr-a still ‘enigmatic multitasker’. Frontiers in Microbiology vol. 5 127 (2014).

27. Gonzalez, M. E. The HIV-1 vpr protein: A multifaceted target for therapeutic intervention. International Journal of Molecular Sciences vol. 18 126 (2017).

28. Lubow, J. & Collins, K. L. Vpr Is a VIP: HIV Vpr and infected macrophages promote viral pathogenesis. Viruses vol. 12 809 (2020).

29. Felzien, L. K. et al. HIV transcriptional activation by the accessory protein, Vpr, is mediated by the p300 co-activator. Proc. Natl. Acad. Sci. U. S. A. 95, 5281–5286 (1998).

30. Connor, R. I., Chen, B. K., Choe, S. & Landau, N. R. Vpr is required for efficient replication of human immunodeficiency virus type-1 in mononuclear phagocytes. Virology 206, 935–944 (1995).

31. Romani, B., Baygloo, N. S., Hamidi-Fard, M., Aghasadeghi, M. R. & Allahbakhshi, E. HIV-1 Vpr Protein Induces Proteasomal Degradation of Chromatin-associated Class I HDACs to Overcome Latent Infection of Macrophages. J Biol Chem 291, 2696–2711 (2016).

32. Poon, B. & Chen, I. S. Y. Human Immunodeficiency Virus Type 1 (HIV-1) Vpr Enhances Expression from Unintegrated HIV-1 DNA. J. Virol. 77, 3962–3972 (2003).

33. Poon, B., Chang, M. A. & Chen, I. S. Y. Vpr Is Required for Efficient Nef Expression from Unintegrated Human Immunodeficiency Virus Type 1 DNA. J. Virol. 81, 10515–10523 (2007).

34. Bauby, H. et al. HIV-1 vpr induces widespread transcriptomic changes in CD4+ T cells early postinfection. MBio 12, (2021).

35. Greenwood, E. J. D. et al. Promiscuous Targeting of Cellular Proteins by Vpr Drives Systems-Level Proteomic Remodeling in HIV-1 Infection. Cell Rep 27, 1579–1596 e7 (2019).

36. Mashiba, M., Collins, D. R., Terry, V. H. & Collins, K. L. Vpr overcomes macrophage-specific restriction of HIV-1 Env expression and virion production. Cell Host Microbe 16, 722–735 (2014).

37. Laguette, N. et al. Premature activation of the SLX4 complex by Vpr promotes G2/M arrest and escape from innate immune sensing. Cell 156, 134–145 (2014).

38. Yan, J., Shun, M. C., Zhang, Y., Hao, C. & Skowronski, J. HIV-1 Vpr counteracts HLTF-mediated restriction of HIV-1 infection in T cells. Proc. Natl. Acad. Sci. U. S. A. 116, 9568–9577 (2019).

39. Wu, Y. et al. The DDB1-DCAF1-Vpr-UNG2 crystal structure reveals how HIV-1 Vpr steers human UNG2 toward destruction. Nat. Struct. Mol. Biol. 23, 933–939 (2016).

40. Zhao, L. et al. Vpr counteracts the restriction of LAPTM5 to promote HIV-1 infection in macrophages. Nat. Commun. 12, 1–14 (2021).

41. Vauthier, V. et al. HUSH-mediated HIV silencing is independent of TASOR phosphorylation on threonine 819. Retrovirology 19, (2022).

42. He, J. et al. Human immunodeficiency virus type 1 viral protein R (Vpr) arrests cells in the G2 phase of the cell cycle by inhibiting p34cdc2 activity. J. Virol. 69, 6705–6711 (1995).

43. Di Marzio, P., Choe, S., Ebright, M., Knoblauch, R. & Landau, N. R. Mutational analysis of cell cycle arrest, nuclear localization and virion packaging of human immunodeficiency virus type 1 Vpr. J. Virol. 69, 7909–7916 (1995).

44. Goh, W. et al. HIV-1 Vpr increases viral expression by manipulation of the cell cycle: A mechanism for selection of Vpr in vivo. Nat. Med. 4, 65–71 (1998).

45. Jacquot, G. et al. Localization of HIV-1 Vpr to the nuclear envelope: Impact on Vpr functions and virus replication in macrophages. Retrovirology 4, (2007).

46. Jacquot, G. et al. Characterization of the molecular determinants of primary HIV-1 Vpr proteins: Impact of the Q65R and R77Q substitutions on Vpr functions. PLoS One 4, e7514 (2009).

47. Vodicka, M. A., Koepp, D. M., Silver, P. A. & Emerman, M. HIV-1 Vpr interacts with the nuclear transport pathway to promote macrophage infection. Genes Dev 12, 175–185 (1998).

48. Le Rouzic, E. et al. HIV1 Vpr arrests the cell cycle by recruiting DCAF1/VprBP, a receptor of the Cul4-DDB1 ubiquitin ligase. Cell Cycle 6, 182–188 (2007).

49. Simmons, G. et al. Primary, Syncytium-Inducing Human Immunodeficiency Virus Type 1 Isolates Are Dual-Tropic and Most Can Use Either Lestr or CCR5 as Coreceptors for Virus Entry. Journal of Virology vol. 70 http://jvi.asm.org/ (1996).

50. Belshan, M., Mahnke, L. A. & Ratner, L. Conserved amino acids of the human immunodeficiency virus type 2 Vpx nuclear localization signal are critical for nuclear targeting of the viral preintegration complex in non-dividing cells. Virology 346, 118–126 (2006).

51. Desai, T. M. et al. Fluorescent protein-tagged Vpr dissociates from HIV-1 core after viral fusion and rapidly enters the cell nucleus. Retrovirology 12, 88 (2015).

52. DeHart, J. L. et al. HIV-1 Vpr activates the G2 checkpoint through manipulation of the ubiquitin proteasome system. Virol. J. 4, (2007).

53. Zhou, Y. & Ratner, L. Phosphorylation of Human Immunodeficiency Virus Type 1 Vpr Regulates Cell Cycle Arrest. J. Virol. 74, 6520–6527 (2000).

54. Stivahtis, G. L., Soares, M. A., Vodicka, M. A., Hahn, B. H. & Emerman, M. Conservation and host specificity of Vpr-mediated cell cycle arrest suggest a fundamental role in primate lentivirus evolution and biology. J. Virol. 71, 4331–4338 (1997).

55. Planelles, V. et al. Vpr-induced cell cycle arrest is conserved among primate lentiviruses. J. Virol. 70, 2516–2524 (1996).

56. Subbramanian, R. A. et al. Human immunodeficiency virus type 1 Vpr is a positive regulator of viral transcription and infectivity in primary human macrophages. J. Exp. Med. 187, 1103–1111 (1998).

57. Cohen, E. A. et al. Identification of HIV-1 vpr product and function. J Acquir Immune Defic Syndr 3, 11–18 (1990).

58. Kruize, Z. & Kootstra, N. A. The Role of Macrophages in HIV-1 Persistence and Pathogenesis. Frontiers in Microbiology vol. 10 2828 (2019).

59. Hendricks, C. M., Cordeiro, T., Gomes, A. P. & Stevenson, M. The Interplay of HIV-1 and Macrophages in Viral Persistence. Frontiers in Microbiology vol. 12 (2021).

60. McNicholl, J. M., Smith, D. K., Qari, S. H. & Hodge, T. Host Genes and HIV: The Role of the Chemokine Receptor Gene CCR5 and Its Allele (Δ32 CCR5). Emerging Infectious Diseases vol. 3 261–271 (1997).

61. Veenhuis, R. T. et al. Monocyte-derived macrophages contain persistent latent HIV reservoirs. Nat. Microbiol. 8, 833–844 (2023).

62. Burdick, R. C. et al. HIV-1 uncoats in the nucleus near sites of integration. Proc. Natl. Acad. Sci. U. S. A. 117, 5486–5493 (2020).

63. Francis, A. C., Marin, M., Prellberg, M. J., Palermino-Rowland, K. & Melikyan, G. B. Hiv-1 uncoating and nuclear import precede the completion of reverse transcription in cell lines and in primary macrophages. Viruses 12, 1234 (2020).

64. Li, C., Burdick, R. C., Nagashima, K., Hu, W. S. & Pathak, V. K. HIV-1 cores retain their integrity until minutes before uncoating in the nucleus. Proc. Natl. Acad. Sci. U. S. A. 118, (2021).

65. Marno, K. M. et al. RNA-Associated Early-Stage Antiviral Factor Is a Major Component of Lv2 Restriction. J. Virol. 91, (2017).

66. Gramberg, T., Sunseri, N. & Landau, N. R. Evidence for an Activation Domain at the Amino Terminus of Simian Immunodeficiency Virus Vpx. J. Virol. 84, 1387–1396 (2010).

67. Bobadilla, S., Sunseri, N. & Landau, N. R. Efficient transduction of myeloid cells by an HIV-1-derived lentiviral vector that packages the Vpx accessory protein. Gene Ther. 20, 514–520 (2013).

68. Simmons, G. et al. Primary, syncytium-inducing human immunodeficiency virus type 1 isolates are dual-tropic and most can use either Lestr or CCR5 as coreceptors for virus entry. J. Virol. 70, 8355–8360 (1996).

69. Cheney, K. M. & McKnight, A. Interferon-Alpha Mediates Restriction of Human Immunodeficiency Virus Type-1 Replication in Primary Human Macrophages at an Early Stage of Replication. PLoS One 5, e13521 (2010).

70. Jackman, J. & O’Connor, P. M. Methods for Synchronizing Cells at Specific Stages of the Cell Cycle. Curr. Protoc. Cell Biol. 00, 8.3.1–8.3.20 (1998).

