## Supplemental Figures and Legends (S1-S3) for "HIV-1 Vpr counteracts TASOR restriction to promote infection prior to integration"

**Figure S1**

**a**

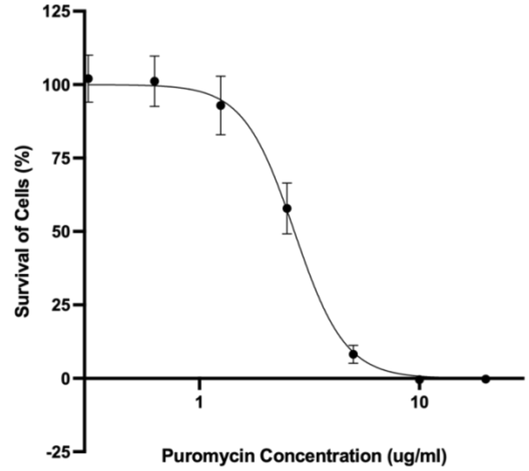
Survival of Cells (%)
125
100
75
50
25
0
-25
1
10
Puromycin Concentration (ug/ml)

**b**

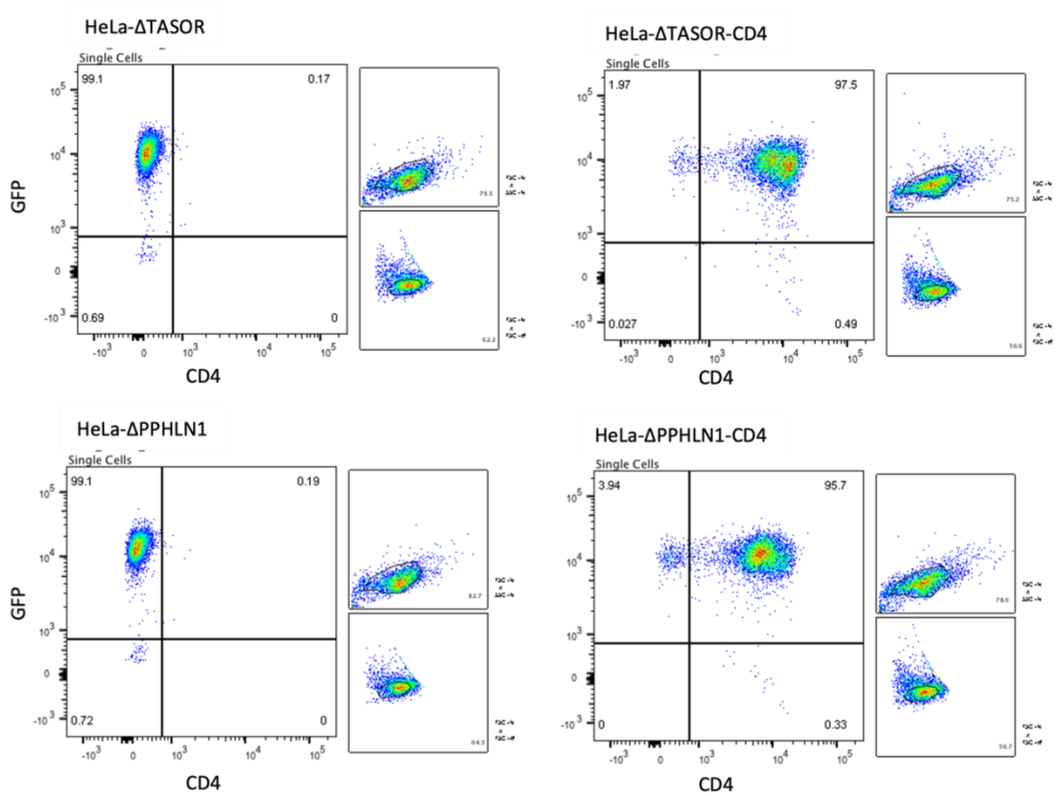
HeLa-ΔTASOR
Single Cells
99.1
0.17
0.69
0
GFP
CD4
HeLa-ΔTASOR-CD4
Single Cells
1.97
97.5
0.027
0.49
CD4
HeLa-ΔPPHLN1
Single Cells
99.1
0.19
0.72
0
GFP
CD4
HeLa-ΔPPHLN1-CD4
Single Cells
3.94
95.7
0
0.33
CD4

**Figure S1**: Puromycin dose-response and flow cytometry of CD4 expression in CRISPR-Cas9 HeLa cells. **a**, Percentage (%) survival of cells treated with different concentrations of puromycin to determine appropriate concentration required for complete kill of non-transfected or transduced cells. **b**, Flow cytometry of hCD4 and GFP in HeLa-ΔTASOR (top, left) and HeLa-ΔPPHLN1 (bottom, left) and in puromycin resistant HeLa-ΔTASOR-CD4 (top, right) and HeLa-ΔPPHLN1-CD4 (bottom, right). Puromycin resistant cell populations expressed high levels of CD4.

**Figure S2**

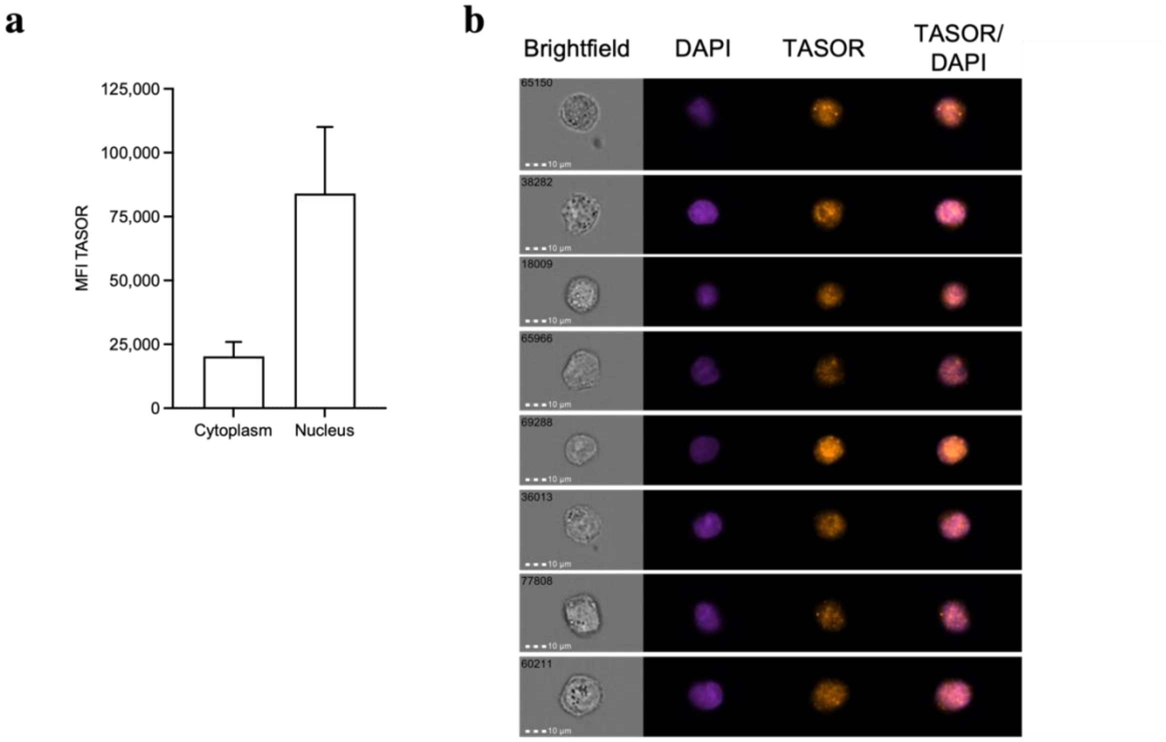
a
b
MFI TASOR
125,000
100,000
75,000
50,000
25,000
0
Cytoplasm
Nucleus
Brightfield
DAPI
TASOR
TASOR/
DAPI
65150
38282
18009
65966
69288
36013
77808
60211
10 μm

**Figure S2**: Imaging flow cytometry analysis of TASOR subcellular localisation. **a**, Imaging flow cytometry of MFI TASOR in the cytoplasmic and nuclear region of HeLa-CD4 cells. Error bars represent standard deviations of the means of technical replicates. **b**, Representative images (60X magnification) of TASOR in the nuclear region of HeLa-CD4 cells. Brightfield, DAPI (purple), TASOR (orange) and merged TASOR/DAPI images are shown. Scale bars 10µm.

**Figure S3**

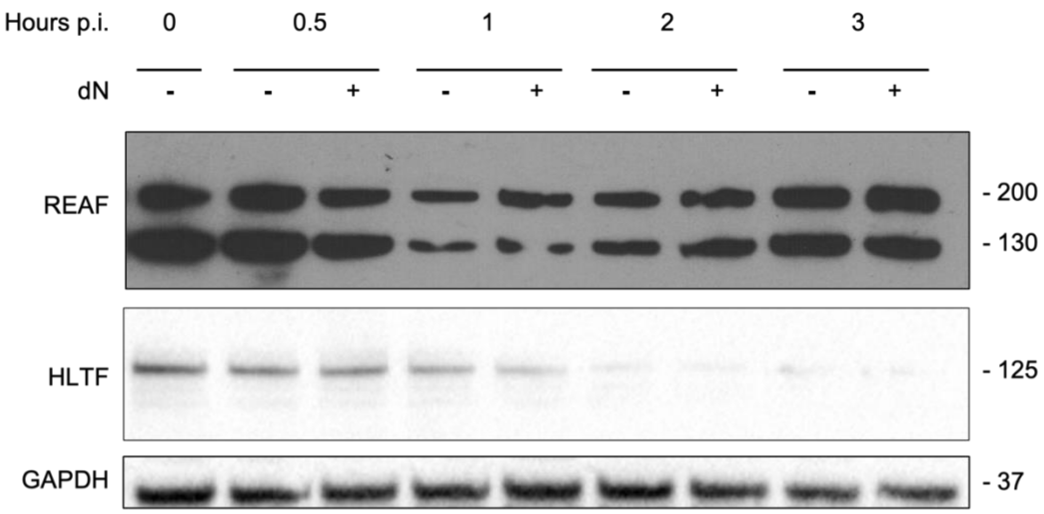

**Figure S3**: Western blotting of REAF and HLTF post infection. Western blotting of REAF and HLTF protein levels in whole cell lysates from THP-1 cells over the first 3 hours post challenge with HIV-1 89.6$^{WT}$. Infections were performed with and without 2'-deoxyneucleoside (dN) supplementation. GAPDH is a control for equal loading.
